# A Calibrated Gate for Evaluating Synthetic Clinical Data

**DOI:** 10.64898/2026.11.22.768435

**Authors:** Neeam Shahriar Hayder, Behraj Khan, Syed Ahmad Chan Bukhari

## Abstract

Evaluation tools for synthetic patient records score fidelity (resemblance to real patients), utility (performance of models trained on the table) and privacy (exposure of real patients), combining them by arithmetic, geometric or harmonic means. Such scores are compensatory: copied training records can pass with high fidelity despite poor privacy. Under the null that a table is an unaltered sample of the source population, none of seven reviewed frame-works reports its aggregation rule’s type I error rate or power against injected flaws. We estimate both for a non-compensatory gate requiring every dimension to pass and for the arithmetic, geometric and harmonic means on identical evidence, each calibrated on the same 50 real-versus-real comparisons by a full-conformal maximum-statistic rule. For exchangeable real tables rejection on complete evidence is bounded by 2/51 (3.9%) at *α* = 0.05, regardless of signal distribution. On 650 tables with injected flaws (13 flaw–dose conditions in each of 50 independent samples from a cohort of 71,518 patients), the arithmetic mean accepted 278 and the gate 170: 2.16 more flawed tables per sample (95% interval 1.56 to 2.76; sign test *p* = 1.0 × 10^−8^). The excess held in all 21 post hoc combinations of score mapping and calibration level (interval excluding zero in 20), with mapping-dependent size; at the prespecified setting, the geometric and harmonic means did not differ detectably from the gate. The fidelity-only SDMetrics quality score accepted 579, including 47 of 50 tables entirely comprising copied training records. With fixed thresholds, the gate rejected or could not evaluate all 50 held-out real-versus-real comparisons; calibrated, it rejected 6 of 200 (3.0%; one-sided 95% upper bound 5.8%). Arithmetic averaging thus admits flaws confined to one dimension; a gate attains a known type I error rate only after calibration on real data. We did not study human reliance on these tools; code and saved results are released.

## 1 Introduction

Before a hospital shares a synthetic copy of its patient records, someone has to decide whether the copy can be trusted. The decision has parts. Does the table look like the patients it came from? Would a model trained on it work on real patients? Could a real patient be found inside it? Each question has well-developed tests [7, 19, 23]. Much less attention has gone to the last step: how the answers are turned into one decision.

That step is not neutral. Take a real table and, in 90% of its rows, shuffle one variable, the number of prior inpatient stays. Every column keeps its distribution. A model trained on the table still predicts readmission almost as well as before, because other variables carry much of the same signal, and no patient is exposed. Only the joint structure is broken, and only a fidelity test notices. Two of the three scores look good; one is poor. An average lets the good scores pay for the poor one, a property the composite-indicator literature calls *compensability* [30, 34]. For clinical data this is the wrong exchange. Accuracy does not repair a distorted cohort, and resemblance does not repair a leaked record.

The obvious remedy is a gate: accept only if every dimension passes. Gates have been proposed for synthetic clinical data [17], and geometric means have been advocated because they behave like an “and” [3]. But a gate has a cost that, to our knowledge, nobody has measured. Because any one dimension can reject a table, a gate inherits the false alarms of its least specific check. Two samples of real patients never agree exactly, so a check with fixed thresholds eventually flags the difference. A gate built from such checks rejects real data.

The two failure modes have names in the study of human trust in automation. *Misuse* is over-reliance on an automated aid; *disuse* is its underutilisation [35]. Trust is appropriate when reliance matches what the system can actually do [21, 25]. A combining rule that certifies flawed tables can invite misuse. A gate that rejects real patients can invite disuse: practitioners may learn to ignore it, and an ignored gate protects no one. An evaluation earns appropriate trust only if both errors are measured.

We therefore ask a question with an empirical answer. *Holding the evidence fixed, does the combining rule change which synthetic tables are accepted? And can a gate reject flawed tables without rejecting real patients?* We answer both on a public cohort, using tables whose flaws we planted ourselves, so the correct verdict is known, and we replicate the comparison across 50 independently drawn real tables. Our contributions are:

1. A controlled comparison of combining rules on identical evidence that measures both errors at matched honest-data cost, with the base table, not the flawed table, as the unit of replication (Sections 3 and 4).
2. A calibration procedure for gates: candidates are ranked against real-versus-real comparisons with a full-conformal rule, a prespecified admission test decides which dimensions may gate, and the result is validated on 200 held-out honest comparisons and 1,000 reshuffled splits (Section 4).
3. Evidence for both halves of the argument (Section 5). Uncalibrated gates rejected nearly all honest data; the calibrated gate rejected 3.0%. At equal cost, the arithmetic mean accepted 278 of 650 single-flaw tables against the gate’s 170, and more than the gate in 38 of 50 bases. The arithmetic mean’s excess persisted under every score mapping we tried; means dominated by their worst input were not consistently ordered with the gate. SDMetrics, which measures fidelity alone, accepted 579, including copies of training patients.

## 2 Related Work

### What is measured

Reviews of synthetic health data agree on the main families of evidence: resemblance to the source (fidelity), usefulness for a downstream task (utility), and disclosure risk (privacy) [16, 19, 22, 23]. They also agree that the field lacks a standard [23, 44], and a multi-expert consensus advises against the most common similarity-based privacy metrics because they “lack precise interpretation” [37]. The dimensions differ in kind. Fidelity is a two-sample question [1, 27, 41]; utility is a transfer question, which in medicine must include calibration [14, 45, 50]; privacy is an adversarial question [6, 13, 40, 42, 52], distinct from formal guarantees [12, 32]. A table can pass one and fail another, and fairness adds subgroup questions on top [4, 33].

### How it is combined

Tools differ much less in what they measure than in what they do next (Table 1). SDMetrics reports an overall quality score that averages its property scores [11, 36]. TabSynDex is designed as a single-score metric built from component scores [8]. Yan et al. rank generators by a weighted sum of rank-derived scores, with equal weights by default [50]. SynthEval min–max normalises each metric and adds the results without weights [24]. Belgodere et al. combine five trust indices with a weighted geometric mean, chosen because it “embodies an ‘and’ operation interpretation, unlike the arithmetic mean, which implies an ‘or’ interpretation” [3]. Synthcity reports metrics and leaves their combination to the user [39]. Requirements that must each be met appear mainly for privacy: SynQP identifies models that “satisfy both quality and privacy” requirements, using the 0.09 identity-risk threshold it attributes to Health Canada and the European Medicines Agency [20]. A non-compensable five-dimension gate with provisional fixed thresholds was proposed in earlier work [17]. Score-cards and quality frameworks for health data organise the same evidence for human review [31, 44, 51]. None of these frameworks, to our knowledge, tests its combining rule against tables whose verdict is known, or measures how often the rule rejects real data.

**Table 1:** How existing frameworks combine evidence, and whether the combining rule itself is evaluated. “Not reported” means we found no such evaluation in the cited work; it does not mean the rule fails.

| Framework | How dimensions are combined | Rule tested on known flaws | Rejection of real data measured |
|---|---|---|---|
| SDMetrics [11] | Arithmetic mean of property scores | Not reported | Not reported |
| TabSynDex [8] | Single score from components | Not reported | Not reported |
| Yan et al. [50] | Weighted sum of rank scores | Not reported | Not reported |
| SynthEval [24] | Unweighted sum of normalised ranks | Not reported | Not reported |
| Belgodere et al. [3] | Weighted geometric mean ("and") | Not reported | Not reported |
| Synthcity [39] | Left to the user | – | – |
| SynQP [20] | Quality and privacy thresholds | Not reported | Not reported |
| Earlier work [17] | Gate, fixed thresholds | Not reported | Not reported |
| This work | Gate calibrated on real data | Yes (950 tables) | Yes (200 runs) |

### Lessons from elsewhere

The question is old in other fields. Builders of composite indicators note that under additive aggregation “a deficit in one dimension can thus be offset (compensated) by a surplus in another”, and that only a non-compensatory approach “could assure non-compensability” [30, 34]. In machine learning, single-number leaderboards and simple averages across metrics have been criticised on similar grounds [10, 15]. Statistics supplies the mechanism. A rule built on the largest of several deviations responds to one strong signal; a rule that adds deviations responds to many weak ones [18, 28]. Tests built on the maximum are powerful against sparse alternatives [5], and one serious flaw is a sparse alternative by construction. A gate is a maximum-type rule; an average is a sum-type rule.

### Calibrating against real data

Judging a synthetic table against a held-out real table, rather than against an ideal of zero difference, is established in fidelity and privacy evaluation [1, 38]. Controlling the error of a decision that takes the maximum over many checks is the classical maxT problem [47]. Conformal inference provides finite-sample control from exchangeability alone [2, 26, 46]. We combine these ideas into a calibrated gate, and then use it to test the combining rule.

## 3 Two Principles

Each gating dimension *k* ∈ *K* = {fid, util, priv} yields a statistic *M* ^(*k*)^(*X*), larger meaning worse (Section 4). A gate accepts a candidate *X* only if every *M* ^(*k*)^(*X*) is at most a threshold. A compensatory rule first maps each statistic to a score *s*_*k*_ ∈ (0, 1] and accepts if the arithmetic (AM), geometric (GM) or harmonic (HM) mean of the *K* = 3 scores is at least a threshold *τ*.

### Principle 1: non-compensation

*A release check should decide each dimension on its own*. The means do not. With *τ* = 0.8 and two dimensions scoring 1, the arithmetic, geometric and harmonic means accept the third at scores as low as 3*τ* − 2 = 0.40, *τ* ^3^ = 0.51 and *τ* /(3 − 2*τ*) = 0.57, all well below *τ* ; whether a gate accepts that dimension does not depend on the others. The means differ in how far they compensate: min_*k*_ *s*_*k*_ ≤ HM ≤ GM ≤ AM, and HM ≤ *K* min_*k*_ *s*_*k*_, so the harmonic mean is dominated by its worst input. Non-compensation is a requirement we choose for clinical data, where a high score on one dimension does not repair a failure on another.

### Principle 2: calibration

*A check may decide a verdict only once its probability of failing a real table is bounded*. This has a price, because a gate fails a table whenever one check does. If check *k* fails a real table with probability *α*_*k*_, the gate’s failure event is the union of the checks’ events, so its probability *α*_*G*_ satisfies max_*k*_ *α*_*k*_ ≤ *α*_*G*_ ≤ ∑_*k*_ *α*_*k*_ whatever the dependence between checks. One check that fails most real tables therefore makes the gate fail most real tables, and calibrating each check separately at level *α* bounds the gate only by *K α*. Section 4 instead calibrates the gate as a whole, through max_*k*_ *M* ^(*k*)^.

### Expectations recorded before scoring

A flaw is *sparse* if it shifts one dimension’s statistic and *dense* if it shifts several slightly. Rules that reject on a maximum are typically more powerful against sparse alternatives, and rules that add evidence against dense ones [5, 28]. The gate rejects on max_*k*_ *M* ^(*k*)^, whereas the geometric mean adds clipped statistics (Section 4). These are heuristics, not theorems for our dependent, clipped signals, so we recorded three expectations before the data they concern were scored: (E1) at a common nominal level, the gate accepts fewer tables with one serious flaw than the arithmetic mean; (E2) the harmonic mean comes close to the gate; (E3) when mild flaws are spread over several dimensions, the gate’s advantage shrinks.

Principle 1 guards against misuse and Principle 2 against disuse [35].

## 4 Method

Table 2 collects the notation used across sections; other symbols are defined where they appear. We fix the data and the null hypothesis, define the evidence and the decision rules, prove the bound that links them, and then define the injected flaws and the analysis.

**Table 2:**
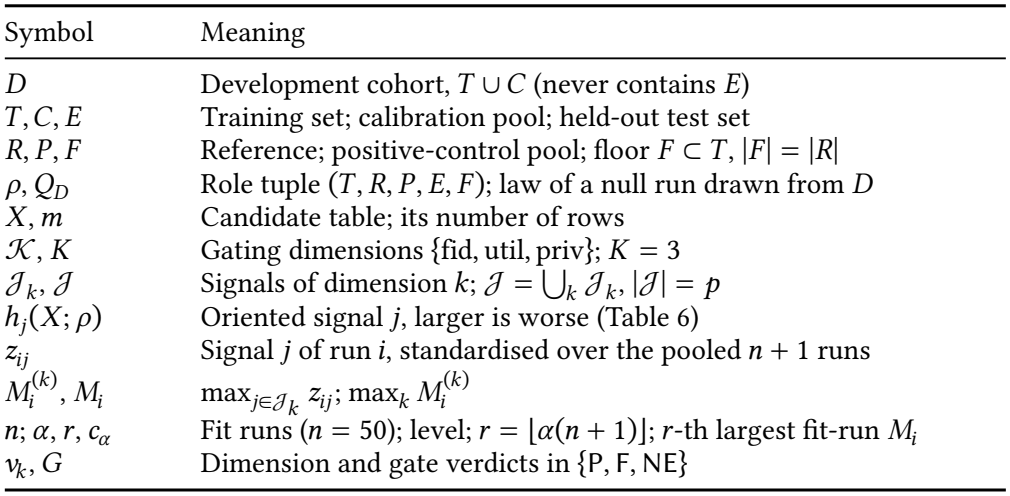
Notation used across sections.

| Symbol | Meaning |
|---|---|
| *D* | Development cohort, *T* ∪ *C* (never contains *E*) |
| *T*, *C*, *E* | Training set; calibration pool; held-out test set |
| *R*, *P*, *F* | Reference; positive-control pool; floor *F* ⊂ *T*, \|*F*\| = \|*R*\| |
| *ρ*, *Q*<sub>*D*</sub> | Role tuple (*T*, *R*, *P*, *E*, *F*); law of a null run drawn from *D* |
| *X*, *m* | Candidate table; its number of rows |
| 𝒦, *K* | Gating dimensions {fid, util, priv}; *K* = 3 |
| 𝒥<sub>*k*</sub>, 𝒥 | Signals of dimension *k*; 𝒥 = ⋃<sub>*k*</sub> 𝒥<sub>*k*</sub>, \|𝒥\| = *p* |
| *h*<sub>*j*</sub>(*X*; *ρ*) | Oriented signal *j*, larger is worse (Table 6) |
| *z*<sub>*ij*</sub> | Signal *j* of run *i*, standardised over the pooled *n* + 1 runs |
| *M*<sub>*i*</sub><sup>(*k*)</sup>, *M*<sub>*i*</sub> | max<sub>*j*∈𝒥<sub>*k*</sub></sub> *z*<sub>*ij*</sub>; max<sub>*k*</sub> *M*<sub>*i*</sub><sup>(*k*)</sup> |
| *n*; *α*, *r*, *c*<sub>*α*</sub> | Fit runs (*n* = 50); level; *r* = ⌊*α*(*n* + 1)⌋; *r*-th largest fit-run *M*<sub>*i*</sub> |
| *ν*<sub>*k*</sub>, *G* | Dimension and gate verdicts in {P, F, NE} |

### Data, roles and the null hypothesis

We use the UCI Diabetes 130-US Hospitals dataset [9, 43], keeping each patient’s earliest encounter (71,518 patients; 30-day readmission prevalence 8.8%); each row has 8 count and 12 categorical variables. Patients are partitioned, stratified by outcome and verified disjoint on hashed identifiers, into a training set *T* (28,607), a calibration pool *C* (28,607) and a test set *E* (14,304); *C* is split into a reference *R* and a positive-control pool *P*, and the floor *F* is a random subset of *T* with |*F* | = |*R*|. In the main roles every compared table has *m* = 14, 303 rows. Every probability statement below concerns a *null run*: a role tuple *ρ* = (*T, R, P, E, F*) drawn from the development cohort *D* = *T* ∪ *C* by a fixed stratified random partition, together with the real candidate *X* = *P*. We write (*X, ρ*) ∼ *Q*_*D*_ for its law, which is conditional on *D*. The null hypothesis for a candidate *X* with roles *ρ* is *H*_0_: (*X, ρ*) *is exchangeable with the null runs*, as holds when (*X, ρ*) ∼ *Q*_*D*_. A rule’s *type I error rate* is Pr{*X* not accepted ∣ *H*_0_}; its *power* against a perturbation is the probability of not accepting a table to which that perturbation was applied. All conclusions are therefore conditional on *D*: they describe the resampling law of this cohort, not new populations.

### Evidence

Each gating dimension *k* ∈ *K* has a fixed set *J*_*k*_ of signals, |*J*_fid_| = 6, |*J*_util_| = 7 and |*J*_priv_| = 3, so *p* = 16. Signal *j* is a real-valued *h*_*j*_ (*X* ; *ρ*), oriented so that larger is worse. Table 6 (Appendix D) defines each one, its reference and its orientation. They are of four kinds: fidelity distances relative to the floor, *ϕ*_*j*_ (*X* ; *R*) − *ϕ*_*j*_ (*F* ; *R*); the exact-match share relative to the real coincidence rate, the share of rows of *X* found in *T* minus the share of rows of *E* found in *T* ; utility quantities relative to a probe trained on *F*, entered through the upper end of a bootstrap interval; and quantities used as computed (the constraint-violation rate, the closer-than-real share and the membership-attack AUC). The evidence of *X* is *complete* if *h*_*j*_ (*X* ; *ρ*) is defined for every *j* ∈ *J*; it is incomplete, for example, when a utility probe trained on real data is not better than chance. Equity and robustness are computed by the same machinery but form a separate diagnostic family that never enters the gate (Section 5).

### Calibration runs

We draw 100 null runs 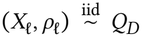 with *m* = 11,443 and |*T*_*l*_| = 22,885; *E* is never read. Patients recur across runs, but each run is an independent random partition of *D*, so the runs are i.i.d. conditional on *D*. Fifty runs form the fit set *ℐ*_fit_ (*n* = 50) and fifty the audit set *ℐ*_aud_; the evidence of every fit run was complete.

### Decision rule

Given a candidate *X* with roles *ρ*, stack the signals of the *n* fit runs and of *X* into *H* ∈ ℝ^(*n*+1)×*p*^, with *H*_*ij*_ = *h*_*j*_ (*X*_*i*_; *ρ*_*i*_) for *i* ≤ *n* and *H*_*n*+1,*j*_ = *h*_*j*_ (*X* ; *ρ*). Standardise each column over all *n* + 1 rows,

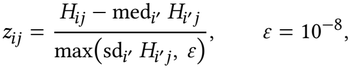

and set 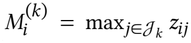 and 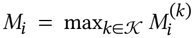. With *r* = ⌊*α*(*n* + 1)⌋, let *c*_*α*_ be the *r* -th largest of *M*_1_, …, *M*_*n*_ (*c*_*α*_ = +∞ if *r* = 0; at *α* ∈ {0.025, 0.05, 0.10}, *r* ∈ {1, 2, 5}). For complete evidence, the verdict of dimension *k* is

- 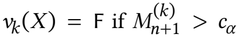, or if *k* = util and *u*(*X*) holds, since no utility probe can be trained on a single outcome class;
- *v*_*k*_ (*X*) = P otherwise.

If the evidence of *X* is incomplete, every dimension with an undefined signal has *v*_*k*_ (*X*) = NE (not evaluated) unless a defined signal of *k* already exceeds *c*_*α*_ ; this is an operational policy, outside the scope of Proposition 4.1. The gate verdict is the most severe dimension verdict under F ≻ NE ≻ P, *G*(*X*) = max_≻_{*v*_*k*_(*X*) : *k* ∈ *K*}, and the gate accepts *X* if and only if *G*(*X*) = P. Because one threshold serves every dimension, 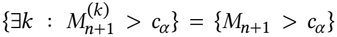, so for complete evidence *G*(*X*) = F if and only if *M*_*n*+1_ > *c*_*α*_ or *u*(*X*): the single-step maxT construction [47] in conformal form.

We also report the conformal *p*-value π (*X*) = (1 + #{*i* ≤ *n* : *M*_*i*_ ≥ *M*_*n*+1_})/(*n* + 1), for which *G*(*X*) = F without *u*(*X*) implies π (*X*) ≤ *r* /(*n* + 1).

### Compensatory rules

The arithmetic, geometric and harmonic means see the same evidence. For every pooled run *i* and dimension *k*, 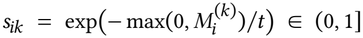 with *t* = 3 fixed in advance, and 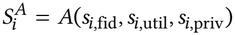 for *A* ∈ {AM, GM, HM}.

Since 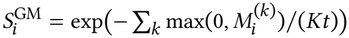 is decreasing in a sum that does not involve *t*, the geometric mean’s verdicts do not depend on *t*. Rule *A* accepts *X* if and only if its evidence is complete, *u*(*X*) does not hold and 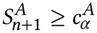, the *r* -th smallest of 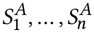.

SDMetrics, an external fidelity-only reference, accepts if and only if its native quality score *q*(*X*) [11] is at least the *r* -th smallest of *q*(*X*_1_), …, *q*(*X*_*n*_); as *q* does not depend on the pooling, this is the split-conformal form of the same rule.

PRoposition 4.1 (Common nominal bound). *Suppose the n* + 1 *pairs* (*X*_1_, *ρ*_1_), *…*, (*X*_*n*_, *ρ*_*n*_), (*X, ρ*) *are exchangeable and every signal is defined for each of them. Then* Pr{*M*_*n*+1_ > *c*_*α*_} ≤ *r* /(*n* + 1), *and the same bound holds for each event* 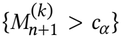, *for*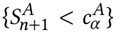 *for each A, and for SDMetrics rejecting. At α* = 0.05 *and n* = 50, *r* /(*n* + 1) = 2/51 = 3.9%.

PRoof. The map *H* ↦ (*M*_1_, …, *M*_*n*+1_) is permutation-equivariant: permuting the rows of *H* permutes the outputs in the same way, because the pooled median and standard deviation are invariant to row order. Hence (*M*_1_, …, *M*_*n*+1_) is exchangeable. Let ℰ_*i*_ be the event that *M*_*i*_ strictly exceeds the *r* -th largest of 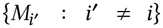. At most *r* events ℰ_*i*_ occur together: if *r* + 1 did, the smallest of those *r* + 1 values would have *r* others at least as large, contradicting the strict inequality. By exchangeability, 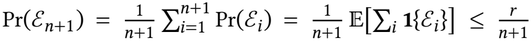, and {*M*_*n*+1_ > *c*_*α*_} = ℰ_*n*+1_. The dimension events are subsets of this event. The same argument applies to −*S*^*A*^, which is also permutation-equivariant, and to −*q*(*X*_*i*_) [26, 46]. □

CoRollaRy 4.2 (Type I eRRoR). *Under H*_0_, *the gate’s type I error rate satisfies* Pr{*G*(*X*) ≠ P} ≤ *r* /(*n*+1)+Pr{*evidence of X incomplete*}+ Pr{*u*(*X*)}, *and the same bound holds for each compensatory rule. For null runs* Pr{*u*(*X*)} = 0, *because roles are stratified by outcome*.

PRoof. *X* is not accepted only if its evidence is incomplete, *u*(*X*) holds, or, with complete evidence, *M*_*n*+1_ > *c*_*α*_ (respectively 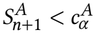; apply the union bound and Proposition 4.1. □

The proposition gives the four rules a *common nominal bound*, not equal error rates: ties and the discrete order statistics can make one rule more conservative than another, so empirical rates are reported separately. The bound is also marginal over the draw of the fit runs: it does not say that a fixed calibration fails at most *r* /(*n*+1) of future real tables. The probability of incomplete evidence is not bounded by the proposition and is estimated on the audit runs.

### Audit and admission

Given the frozen calibration, the audit runs are i.i.d. draws from *Q*_*D*_, which the binomial calculations below require; exchangeability alone would not suffice. For a check *k* ∈ *K* ∪ {*G*}, let 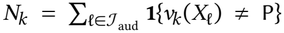 be the number of audit runs it does not accept. Check *k* is *admitted* if and only if *v*_*k*_ (*X*_*l*_) ≠ NE for all *l* ∈ *ℐ*_aud_ and Pr{*B* ≥ *N*_*k*_} ≥ 0.05 for *B* ∼ Binomial(50, 0.05). This one-sided exact test detects gross miscalibration but cannot certify a rate near 5%: it admits *N*_*k*_ = 5 (10%; *p* = 0.10) and excludes *N*_*k*_ = 6 (*p* = 0.04). We therefore also report the exact one-sided 95% (Clopper–Pearson) upper bound on each admitted check’s rate. Admission never changes *c*_*α*_ : the gate family *K* was fixed before the audit, and a check that fails admission is reported, not re-tuned, so Proposition 4.1 still applies to the reported gate.

### Bases and perturbations

A base is the real candidate *X*_*b*_ = *P*_*b*_ of audit run *b* ∈ *ℐ*_aud_, scored against its own roles *ρ*_*b*_, so *m* = 11,443. Unflawed, it is a draw from *Q*_*D*_ and satisfies *H*_0_ by construction. A perturbation family *f* acts through an operator Ψ_*f*_ (*X*_*b*_ ; *d, ω*_*b,f*_) that keeps the size and columns of *X*_*b*_, at dose *d* with one random stream *ω*_*b,f*_ per base and family. Let Δ_*f*_ (*d*) be the set of cells the operator targets. The stream is used so that the targeted rows, the donor rows and the transformation applied to each earlier block are retained when the dose increases; under this coupling, *d* ≤ *d*^′^ ⇒ Δ_*f*_ (*d*) ⊆ Δ_*f*_ (*d*^′^). A targeted cell may keep its value, for example when a rotation moves equal values, so the cells whose values actually change form a subset of Δ_*f*_ (*d*).

- *Copied rows* (*d* ∈ {0.1, 1, 5}%): ⌈*d m*⌉ rows chosen uniformly are overwritten by rows drawn uniformly, with replacement, from *T*_*b*_.
- *Shuffled predictor* (*d* ∈ {10, 50, 90}%): the values of number_inpatient are rotated within disjoint blocks covering ⌈*d m*⌉ rows, the blocks of a lower dose being contained in those of a higher one; the column’s marginal distribution in *X*_*b*_ is unchanged.
- *Missing values* (*d* ∈ {5, 15, 30}%): in each of time_in_hospital, num_lab_procedures, num_medications and number_inpatient, ⌈*d* ⋅ *o*_*c*_ ⌉ of its *o*_*c*_ observed values are set missing, chosen uniformly.
- *Suppressed subgroup* (*d* ∈ {10, 50, 90}%): of the African-American rows *G*, ⌈*d* |*G*|⌉ are replaced by rows drawn with replacement from the other rows of the table.
- *Copy control*: *X* is a sample of *m* rows of *T*_*b*_ drawn without replacement.

These are *prespecified perturbation alternatives*. Each is designed to move *X*_*b*_ away from *H*_0_ in a way a release check exists to detect, and before any confirmatory job ran each family was assigned the gating dimensions *κ*(*f*) ⊆ *K* it targets: copied rows and the copy control, privacy; suppressed subgroup, fidelity; shuffled predictor and missing values, fidelity and utility. That a realised table departs from *H*_0_ is a property of the cohort, not of the operator: shuffling a variable independent of all others, for instance, would leave the joint distribution unchanged. We therefore call the perturbed tables *known-bad* only in this operational sense, which is a statement about the evidence and not a claim of clinical harm; every family and dose is reported separately (Table 4). The 12 family–dose conditions and the copy control form a set *C* of 13 conditions and the primary known-bad set of 13 × 50 = 650 tables. *Negative controls*, scored in the main roles, permute every column of a real table independently, which preserves each marginal and targets every dependence between columns; they are expected to fail fidelity and utility.

### Mild perturbations over several dimensions

To probe the dense case, three mild components were fixed before scoring, one per gating dimension: F, the 50% shuffled predictor (fidelity); U, the outcome rotated among 10% of rows, which leaves prevalence unchanged (utility); and P, one row overwritten by a row of *T*_*b*_ (privacy). Components are applied in the order F, U, P, each with its own stream. A combination targets at most the union of its components’ cells, Δ_F+U+P_ ⊆ Δ_F_ ∪ Δ_U_ ∪ Δ_P_, and not always all of it, because P overwrites a whole row and can replace cells that F or U changed. We score U, P, F+U, F+P, U+P and F+U+P on every base (300 tables); F alone is the 50% shuffle above. The same single perturbations are applied to Gaussian Copula tables as a secondary set, and four generators are fitted with three seeds each: Gaussian Copula, CTGAN and TVAE (SDV 1.37.4, 300 epochs) [36, 49] and AIM with *ε*_DP_ = 1 [29].

### Statistical analysis

Perturbed tables built on one base share it, so the base is the unit of replication. Write *X*_*b,c*_ for base *b* under condition *c* ∈ *C*, and *a*_*J*_ (*X*) ∈ {0, 1} for acceptance by rule *J*, with *a*_*G*_ (*X*) = **1**{*G*(*X*) = P}. For each rule *J* and base *b*,

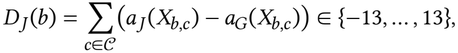

so *D*_*J*_ (*b*) > 0 means that *J* accepted more of base *b*’s known-bad tables than the gate; the 13 conditions are weighted equally. We report 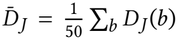 with an approximate 95% bootstrap interval over bases (10,000 resamples), and an exact two-sided sign test of Pr{*D*_*J*_ (*b*) > 0} = Pr{*D*_*J*_ (*b*) < 0} over bases with *D*_*J*_ (*b*) ≠ 0, which concerns the direction of the differences, not their mean. An interval that contains zero is reported as no detectable difference. Per-condition acceptance counts *A*_*J, c*_ = ∑_*b*_ *a*_*J*_ (*X*_*b,c*_) carry Wilson intervals [48]. All of these are conditional on *D* and on the frozen calibration.

### Prespecification

The protocol was amended twice, each time before the data it governs were scored, and every change is logged with its reason. The first amendment followed a 40-run pilot, gated on all five dimensions, whose audit showed that the robustness rule fails real tables (15 of 20), that equity is often not evaluable, and that the earlier calibration under-covered through one zero-inflated signal; the pilot is reported in the released files. The second followed review, which noted that the original known-bad set rested on one base table that was larger than the calibration runs. It fixed the 50-base design, the dense components, the analyses above and how each outcome would be interpreted, and it added 150 supplementary null runs, drawn i.i.d. from *Q*_*D*_ with new seeds, as a further audit. The frozen calibration model, evaluators and judges were reused unchanged, with their code and model verified by hash.

## 5 Results

### 5.1 An uncalibrated gate rejects real patients

Table 3 gives, for each dimension and three versions of the gate, how many of the 50 held-out honest comparisons were not accepted. The frozen v0.3 gate, with fixed thresholds, failed or refused all of them (50/50). Its fidelity check alone rejected every run, and the same gate failed all three positive controls. Measuring signals relative to the real floor, but without calibration, still left 41/50 unaccepted, mostly through privacy (32/50) and utility (20/50). After calibration, the gate rejected 2/50 (4%; 95% CI 1.1–13.5%), against a nominal 3.9%. Fidelity, utility and privacy each passed the admission rule. Two dimensions often listed among the pillars of trust did not. Equity could not be evaluated on 48 of the 50 runs, because some subgroup always lacked a reliable real-data baseline. Robustness rejected every run. We report both, but they do not gate. In the main roles, the calibrated gate rejected every negative control and all 12 generator tables, each of which failed fidelity and utility (Appendix A).

**Table 3:**
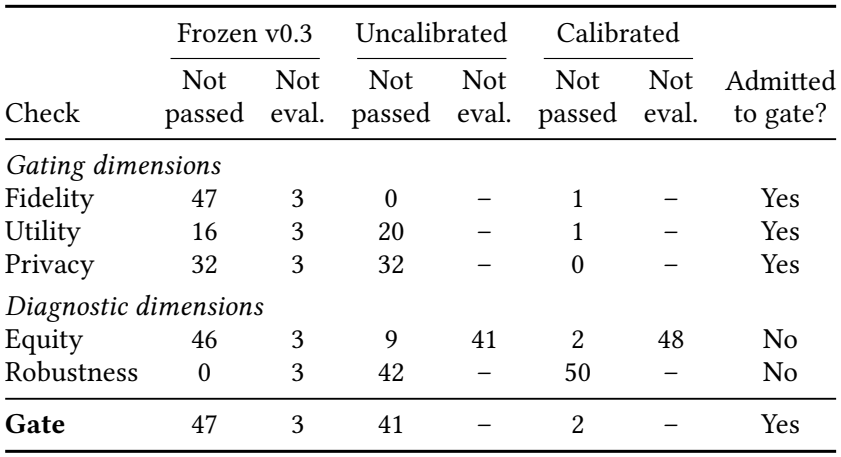
False alarms on honest data: of 50 held-out realversus-real comparisons, how many each check did not pass, and how many it could not evaluate. Every honest comparison should pass, so lower is better; the calibrated rule’s nominal false-alarm rate is about 2 in 50. “Admitted” applies the prespecified admission rule to the calibrated version.

|  | Frozen v0.3 |  | Uncalibrated |  | Calibrated |  |  |
|---|---|---|---|---|---|---|---|
| Check | Not passed | Not eval. | Not passed | Not eval. | Not passed | Not eval. | Admitted to gate? |
| *Gating dimensions* |  |  |  |  |  |  |  |
| Fidelity | 47 | 3 | 0 | – | 1 | – | Yes |
| Utility | 16 | 3 | 20 | – | 1 | – | Yes |
| Privacy | 32 | 3 | 32 | – | 0 | – | Yes |
| *Diagnostic dimensions* |  |  |  |  |  |  |  |
| Equity | 46 | 3 | 9 | 41 | 2 | 48 | No |
| Robustness | 0 | 3 | 42 | – | 50 | – | No |
| **Gate** | 47 | 3 | 41 | – | 2 | – | Yes |

**Table 4:**
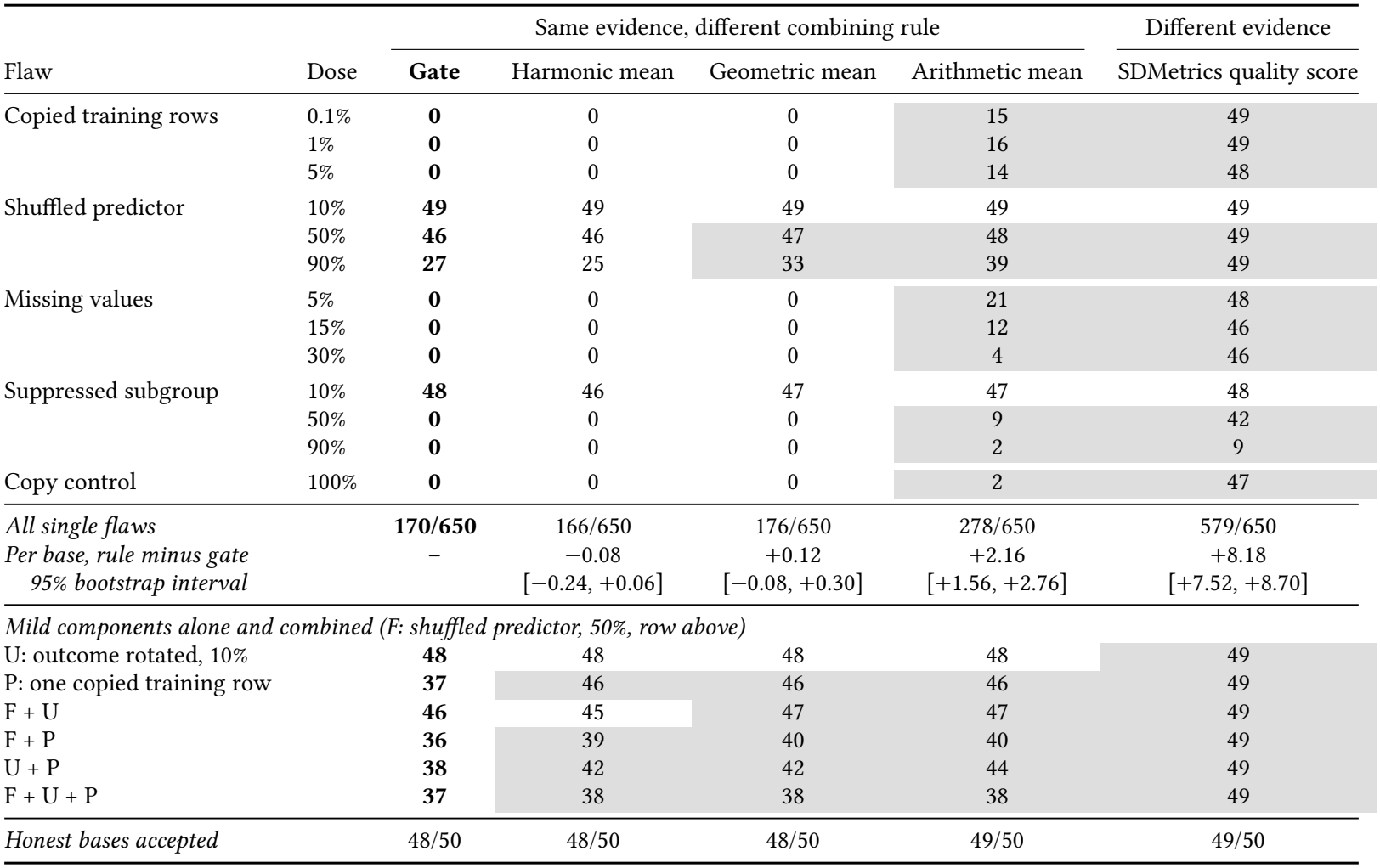
Flawed tables wrongly accepted at matched honest-data cost (*α* = 0.05), of 50 independent bases. Each flaw is injected into a held-out real table; under our acceptance requirement (only an unaltered sample of the source population should pass), the correct verdict for every row is rejection, which for the mildest doses is a strict standard. Lower is better. Shaded cells mark a rule that accepted more of the 50 tables than the gate; differences of one or two tables are within noise, which is why the per-base comparison (middle rows) is the primary analysis. SDMetrics is shown separately because it scores fidelity only and therefore sees different evidence.

| Flaw | Dose | Same evidence, different combining rule |  |  |  | Different evidence |
|---|---|---|---|---|---|---|
|  |  | **Gate** | Harmonic mean | Geometric mean | Arithmetic mean | SDMetrics quality score |
| Copied training rows | 0.1% | **0** | 0 | 0 | 15 | 49 |
|  | 1% | **0** | 0 | 0 | 16 | 49 |
|  | 5% | **0** | 0 | 0 | 14 | 48 |
| Shuffled predictor | 10% | **49** | 49 | 49 | 49 | 49 |
|  | 50% | **46** | 46 | 47 | 48 | 49 |
|  | 90% | **27** | 25 | 33 | 39 | 49 |
| Missing values | 5% | **0** | 0 | 0 | 21 | 48 |
|  | 15% | **0** | 0 | 0 | 12 | 46 |
|  | 30% | **0** | 0 | 0 | 4 | 46 |
| Suppressed subgroup | 10% | **48** | 46 | 47 | 47 | 48 |
|  | 50% | **0** | 0 | 0 | 9 | 42 |
|  | 90% | **0** | 0 | 0 | 2 | 9 |
| Copy control | 100% | **0** | 0 | 0 | 2 | 47 |
| *All single flaws* |  | **170/650** | 166/650 | 176/650 | 278/650 | 579/650 |
| *Per base, rule minus gate* |  | – | −0.08 | +0.12 | +2.16 | +8.18 |
| *95% bootstrap interval* |  |  | [−0.24, +0.06] | [−0.08, +0.30] | [+1.56, +2.76] | [+7.52, +8.70] |
| *Mild components alone and combined (F: shuffled predictor, 50%, row above)* |  |  |  |  |  |  |
| U: outcome rotated, 10% |  | **48** | 48 | 48 | 48 | 49 |
| P: one copied training row |  | **37** | 46 | 46 | 46 | 49 |
| F + U |  | **46** | 45 | 47 | 47 | 49 |
| F + P |  | **36** | 39 | 40 | 40 | 49 |
| U + P |  | **38** | 42 | 42 | 44 | 49 |
| F + U + P |  | **37** | 38 | 38 | 38 | 49 |
| *Honest bases accepted* |  | 48/50 | 48/50 | 48/50 | 49/50 | 49/50 |

Fifty audit runs can rule out gross miscalibration, not a small excess over the nominal rate (upper bound 12.1%). Two further checks tighten this. The 150 fresh honest runs were rejected 4/150 times. Pooled with the audit, the gate rejected 6/200 (3.0%; one-sided 95% upper bound 5.8%): 2 through fidelity, 4 through utility and none through privacy. Reshuffling the 100 original runs into 1,000 random fit/audit splits and refitting gave a mean gate rejection rate of 3.92%, against the nominal 3.92% (Appendix C). The calibration holds on runs it never saw, not only on one split.

### 5.2 At equal cost, averaging hides failures

Figure 2b and Table 4 give the primary comparison. With every rule calibrated to the same honest-data rejection rate, the gate accepted 170 of the 650 single-flaw tables (26%), the harmonic mean 166, the geometric mean 176, the arithmetic mean 278 (43%) and SDMetrics 579 (89%). Acceptance of the unflawed bases was nearly identical (48 to 49 of 50), so these differences were not bought by treating real data more harshly. Per base, the arithmetic mean accepted on average +2.16 flawed tables relative to the gate (95% interval +1.56 to +2.76). It accepted more in 38 of the 50 bases and fewer in 3 (sign test *p* = 1.0 × 10^−8^). For SDMetrics the difference was +8.18 (+7.52 to +8.70; more in 49 bases). The gap held at the other calibration levels: at *α* = 0.025 and 0.10 the gate accepted 182 and 148 single-flaw tables, the arithmetic mean 298 and 213.

**Figure 1:**
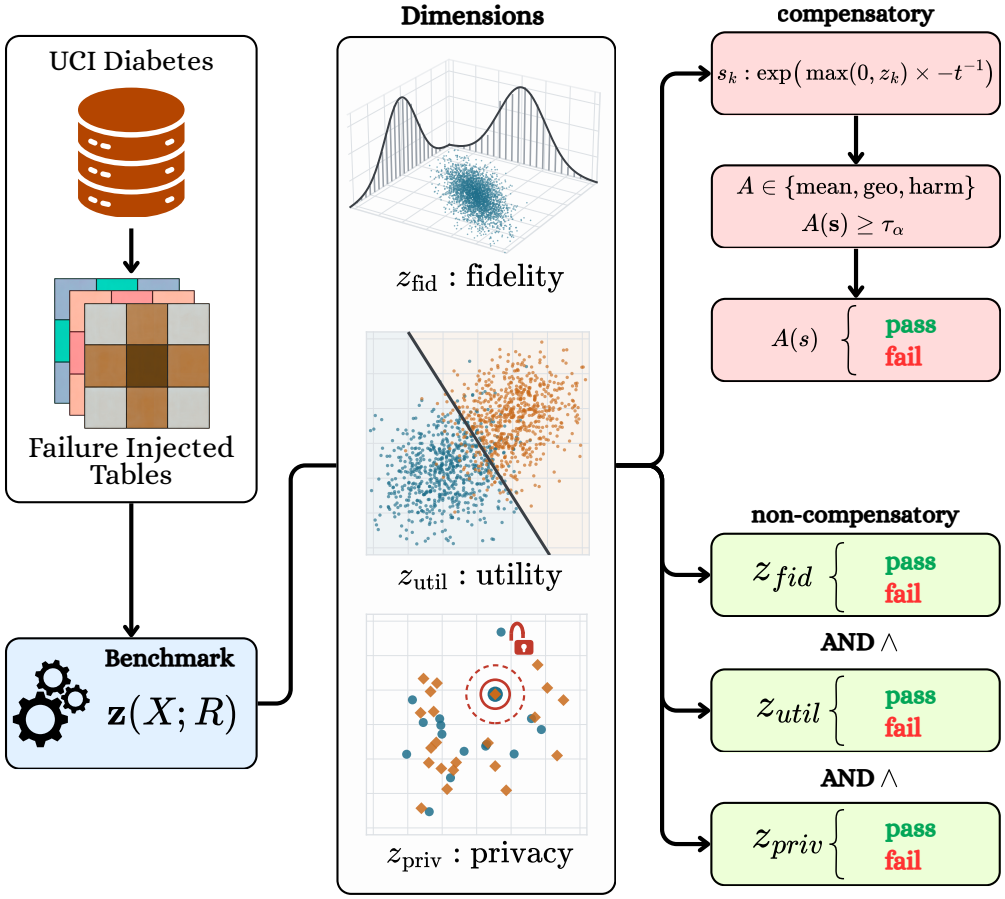
Study design. Each flawed real table *X* is scored against a real reference *R*, and the same evidence z(*X* ; *R*) feeds a compensatory rule (average of scores *s*_*k*_, accept if ≥ *τ*_*α*_) and a gate (accept only if every dimension passes *c*_*α*_), with all thresholds set on separate real-versus-real runs at the same rate *α*. Colours: outcome classes (utility); training versus released rows (privacy).

**Figure 2:**
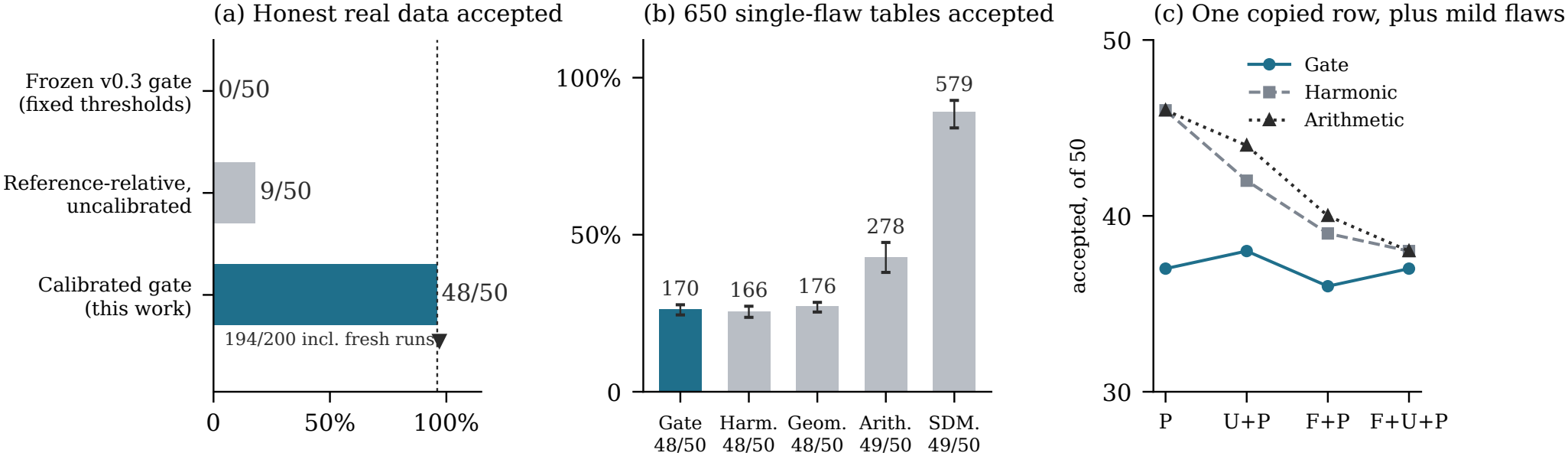
The two errors of a combining rule. (a) Held-out honest real-versus-real comparisons accepted by three versions of the gate; the dashed line is the calibrated rule’s nominal acceptance rate, 49/51, and the triangle adds 150 fresh honest runs. (b) The 650 single-flaw tables (13 conditions on 50 bases) accepted by the gate and the three means, which share the same evidence, and, for reference, by the fidelity-only SDMetrics score. All are calibrated to the same honest-data rejection rate (*α* = 0.05); each rule’s acceptance of the 50 unflawed bases is shown under its name. Bars show 95% bootstrap intervals over bases. (c) Tables with one copied training row (P) and mild fidelity (F) and utility (U) flaws added, accepted of 50; the axis starts at 30.

Against the means dominated by their lowest input, there was no detectable difference. Per base, the harmonic mean differed from the gate by −0.08 tables (−0.24 to +0.06) and the geometric mean by +0.12 (−0.08 to +0.30). We had predicted that the harmonic mean would come close to the gate; it came all the way. Neither rule is uniformly stricter than the other, and the order depends on the calibration level (Section 5.6). Across all single-flaw tables, the gate accepted 10 that the harmonic mean rejected and the harmonic mean 6 that the gate rejected, and the totals swap order between *α* = 0.025 (gate 182, harmonic 170) and *α* = 0.10 (148 and 159).

Table 4 shows where the hidden failures come from. Missing values and the stronger subgroup suppressions were rejected in every base by the gate and by the geometric and harmonic means, while the arithmetic mean accepted up to 21 of 50. Copied training rows were rejected in every base by every rule that weighs privacy hard, while the arithmetic mean accepted 14 to 16 of 50 at each dose. SDMetrics accepted nearly every flawed table, including 47 of 50 copy controls. A copy of real patients is a perfect likeness, and a quality score that measures only likeness rates it highly.

### 5.3 When harm is spread out

A maximum-type rule is expected to lose its edge when harm is spread thinly over several dimensions [28]. Figure 2c and the lower part of Table 4 show what happened. On their own, the fidelity and utility components were nearly invisible: every rule accepted F in at least 46 of 50 bases and U in 48. The single copied row was detected more often, but not reliably. The gate rejected it in 13 of 50 bases and missed it in 37 (74%); each mean rejected it in 4. One copied record barely exceeds the matches that two real samples share by chance, so the privacy check fires only sometimes, and when it does, a mean averages it with two healthy dimensions. Adding the mild components moved the means toward the gate. The harmonic mean accepted 46, 42, 39 and 38 of P, U+P, F+P and F+U+P, while the gate stayed between 36 and 38. The gate’s advantage shrank as components were added, as predicted, but did not reverse. Over the 300 tables in this block, the harmonic mean accepted 21 that the gate rejected, and the gate 5 that the harmonic mean rejected; this paired count was not prespecified and is descriptive. Because F and U were too mild for any rule to detect, the test speaks mainly to a sparse privacy flaw with small additions, not to dense harm in general.

### 5.4 Anatomy of a hidden failure

Why does averaging fail on exactly these tables? Where the rules disagree, one dimension collapses and the others pass. With 5% of values missing, the median fidelity score *s*_*k*_ (1 means no worse than the median honest run) was 0.01, and fidelity passed in none of the 50 bases, while utility (0.63) and privacy (0.74) passed in 49 and 50. With 0.1% copied rows, privacy fell below 0.01 in every base while fidelity and utility stayed near their honest medians (0.64 and 0.66 against 0.65). This is the sparse alternative of Section 3, made concrete: of the 480 single-flaw tables the gate rejected, 394 failed exactly one gating dimension, 79 failed two and 7 all three. An average spreads one collapse over three dimensions; a gate reports it.

### 5.5 What no tested rule could see

The flaws that every rule accepted were the weakest: 10% shuffling (49 of 50 bases) and 10% subgroup suppression (46 to 48). At these doses the evidence itself did not separate the flawed table from real data, and none of the tested rules recovered the flaw. The 90% shuffle lies at the edge: its fidelity check passed in 27 of 50 bases, and there the rules disagreed in both directions. The gate’s limit is the evidence’s limit, not the combination’s. Detecting a 10% suppression of one subgroup probably needs a dimension built for subgroups, and that is the dimension that failed admission.

### 5.6 Sensitivity of the comparison

The means depend on how evidence becomes a score, so we recomputed the matched comparison from saved evidence, post hoc, for five temperatures *t*, a percentile mapping and a model without the closer-than-real signal, at all three levels (Table 5, Appendix B). The arithmetic mean accepted more single-flaw tables than the gate in all 21 settings (95% interval excluding zero in 20), but the mapping largely set the size of its excess: at *α* = 0.05 it fell from +5.28 tables per base at *t* = 1 to +0.30 at *t* = 10. The geometric mean, whose verdicts do not depend on *t*, matched the gate except at *α* = 0.10 (+0.34). The harmonic mean’s direction depended on *α*, every mean accepted far more than the gate under the percentile mapping, and removing the closer-than-real signal left the pattern unchanged. What is robust is qualitative: arithmetic averaging accepted more flawed tables than the gate under every mapping, and the worst-dominated means and the gate were not reliably ordered.

**Table 5:**
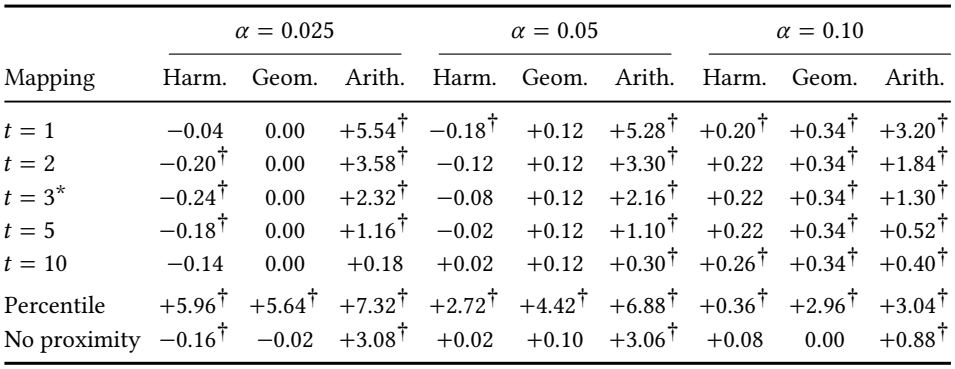
Sensitivity of the matched comparison on the 50 bases (post hoc, from saved evidence). Each cell is the mean per-base difference *D*_*J*_ over the 13 single-flaw conditions: tables accepted by mean *J* minus tables accepted by the gate (positive means the gate accepted fewer). ^†^95% bootstrap interval over bases excludes zero. The gate does not use scores; it accepted 182, 170 and 148 of 650 at the three levels in every row. ^∗^Prespecified. Percentile: each dimension scored by its rank among the pooled runs. No proximity: closer-than-real signal removed. Full intervals and counts are in the released analysis files.

|  | *α* = 0.025 |  |  | *α* = 0.05 |  |  | *α* = 0.10 |  |  |
|---|---|---|---|---|---|---|---|---|---|
| Mapping | Harm. | Geom. | Arith. | Harm. | Geom. | Arith. | Harm. | Geom. | Arith. |
| *t* = 1 | −0.04 | 0.00 | +5.54<sup>†</sup> | −0.18<sup>†</sup> | +0.12 | +5.28<sup>†</sup> | +0.20<sup>†</sup> | +0.34<sup>†</sup> | +3.20<sup>†</sup> |
| *t* = 2 | −0.20<sup>†</sup> | 0.00 | +3.58<sup>†</sup> | −0.12 | +0.12 | +3.30<sup>†</sup> | +0.22 | +0.34<sup>†</sup> | +1.84<sup>†</sup> |
| *t* = 3<sup>\*</sup> | −0.24<sup>†</sup> | 0.00 | +2.32<sup>†</sup> | −0.08 | +0.12 | +2.16<sup>†</sup> | +0.22 | +0.34<sup>†</sup> | +1.30<sup>†</sup> |
| *t* = 5 | −0.18<sup>†</sup> | 0.00 | +1.16<sup>†</sup> | −0.02 | +0.12 | +1.10<sup>†</sup> | +0.22 | +0.34<sup>†</sup> | +0.52<sup>†</sup> |
| *t* = 10 | −0.14 | 0.00 | +0.18 | +0.02 | +0.12 | +0.30<sup>†</sup> | +0.26<sup>†</sup> | +0.34<sup>†</sup> | +0.40<sup>†</sup> |
| Percentile | +5.96<sup>†</sup> | +5.64<sup>†</sup> | +7.32<sup>†</sup> | +2.72<sup>†</sup> | +4.42<sup>†</sup> | +6.88<sup>†</sup> | +0.36<sup>†</sup> | +2.96<sup>†</sup> | +3.04<sup>†</sup> |
| No proximity | −0.16<sup>†</sup> | −0.02 | +3.08<sup>†</sup> | +0.02 | +0.10 | +3.06<sup>†</sup> | +0.08 | 0.00 | +0.88<sup>†</sup> |

**Table 6:**
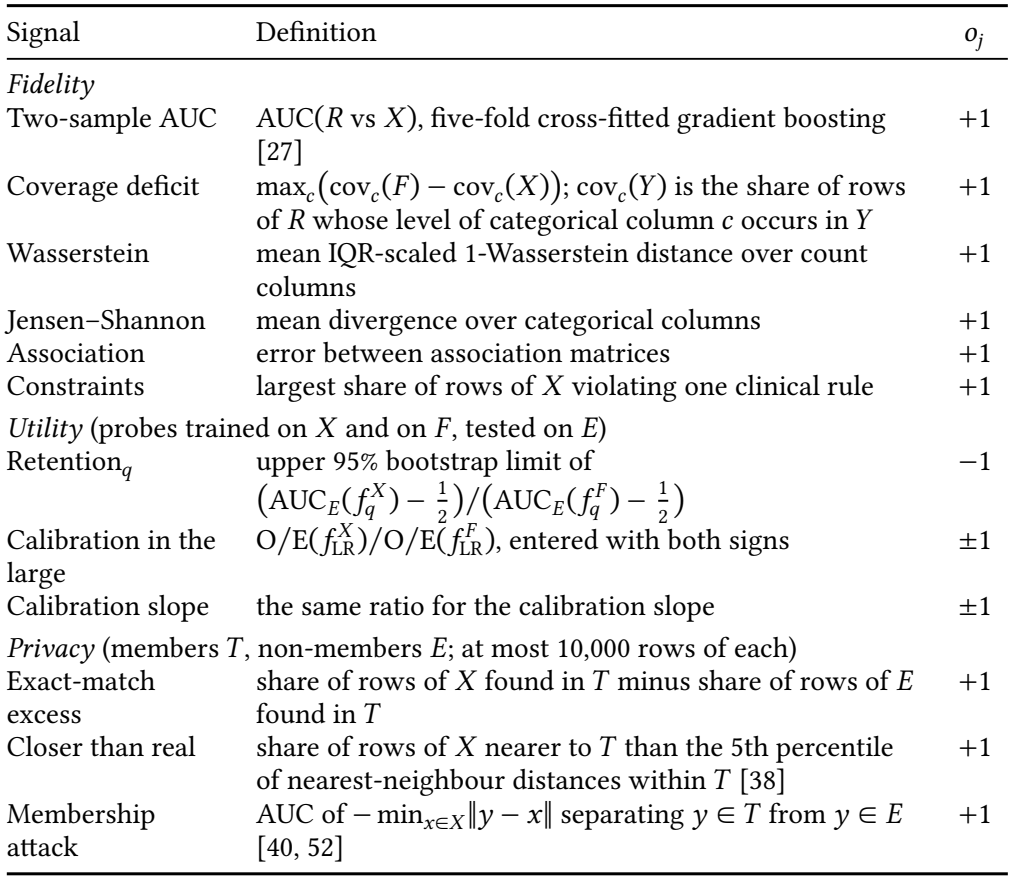
Gating signals, oriented so that larger is worse (*o*_*j*_ is the sign applied). Fidelity quantities are computed against the reference *R* and entered minus the same quantity for the floor *F*, except the constraint rate; privacy quantities other than the exact-match excess are entered as computed.

| Signal | Definition | *o*<sub>*j*</sub> |
|---|---|---|
| *Fidelity* |  |  |
| Two-sample AUC | AUC(*R* vs *X*), five-fold cross-fitted gradient boosting [27] | +1 |
| Coverage deficit | max<sub>*c*</sub>(cov<sub>*c*</sub>(*F*) − cov<sub>*c*</sub>(*X*)); cov<sub>*c*</sub>(*Y*) is the share of rows of *R* whose level of categorical column *c* occurs in *Y* | +1 |
| Wasserstein | mean IQR-scaled 1-Wasserstein distance over count columns | +1 |
| Jensen–Shannon | mean divergence over categorical columns | +1 |
| Association | error between association matrices | +1 |
| Constraints | largest share of rows of *X* violating one clinical rule | +1 |
| *Utility* (probes trained on *X* and on *F*, tested on *E*) |  |  |
| Retention<sub>*q*</sub> | upper 95% bootstrap limit of (AUC<sub>*E*</sub>(*f*<sub>*q*</sub><sup>*X*</sup>) − ½)/(AUC<sub>*E*</sub>(*f*<sub>*q*</sub><sup>*F*</sup>) − ½) | −1 |
| Calibration in the large | O/E(*f*<sub>LR</sub><sup>*X*</sup>)/O/E(*f*<sub>LR</sub><sup>*F*</sup>), entered with both signs | ±1 |
| Calibration slope | the same ratio for the calibration slope | ±1 |
| *Privacy* (members *T*, non-members *E*; at most 10,000 rows of each) |  |  |
| Exact-match excess | share of rows of *X* found in *T* minus share of rows of *E* found in *T* | +1 |
| Closer than real | share of rows of *X* nearer to *T* than the 5th percentile of nearest-neighbour distances within *T* [38] | +1 |
| Membership attack | AUC of − min<sub>*x*∈*X*</sub>‖*y* − *x*‖ separating *y* ∈ *T* from *y* ∈ *E* [40, 52] | +1 |

### 5.7 The original analysis

Our first confirmatory run injected the same flaws into three positive controls from *P*, so its 39 flawed tables rest on one base table, larger (14,303 rows) than the calibration runs. It pointed the same way (gate 9/39, arithmetic mean 17/39, SDMetrics 39/39), but its small edge over the geometric and harmonic means did not replicate across bases. Thresholds set from its positive controls rather than matched on honest data made every mean look stricter only by rejecting most real data.

### 5.8 A caution about privacy evidence

All three AIM tables failed calibrated privacy, although AIM is differentially private at *ε* = 1. Only one of them contained a training record (a single row of 14,303), and the membership attack stayed at chance (AUC 0.50). The failing signal was the share of rows closer to a training record than real data are: 15.4–15.8% for AIM, against a real-versus-real median of 5.0%. Coarse public discretisation places many synthetic rows on common patient profiles, and a distance criterion reads this typicality as risk. This agrees with the consensus warning about similarity-based privacy metrics [37, 42]. Removing the signal from the frozen model confirms the diagnosis: all three AIM tables then pass privacy, but still fail fidelity and utility. On the bases, the gate without it still rejected 2 of 50 honest runs and 4/150 fresh ones, and the matched comparison kept its shape (Section 5.6). The episode shows that a gate is only as meaningful as the evidence it receives.

## 6 Discussion

### What the results mean

A combined score is a decision rule, and it can be tested like one. At matched honest-data cost, arithmetic averaging and a fidelity-only quality score certified flawed real data that a calibrated gate rejected, while the means most dominated by their worst input were not consistently ordered with the gate; the geometric mean largely earns its reputation as an “and” [3]. The dividing line is therefore less gate versus mean than whether a rule lets a good dimension pay for a bad one. What distinguishes the gate is structural: each dimension holds a veto with its own audited error rate, a rejection names the dimension that failed, and no score mapping has to be chosen. Its detection advantage over the worst-dominated means was limited to a single copied training row (13 of 50 bases against 4).

### What strictness costs

The second finding is less familiar and, we think, more important in practice. A gate is safe only once it is calibrated. Strictness alone, with thresholds that sound reasonable, produced a gate that rejected every honest comparison. Such a gate does not protect patients, and it risks teaching its users to stop listening [25, 35]. Calibration turns “strict” into “strict at a known price”, and it makes that price auditable.

### What to do

We suggest three practices to anyone who turns evidence about synthetic patient data into a release decision. First, report a gate verdict alongside any aggregate score, and do not rank tables by an average alone. If a scalar is needed, prefer a rule dominated by the worst dimension, calibrate it, and report its score mapping. Second, calibrate every check that can block a release against real-versus-real comparisons, and audit the result on comparisons held out from calibration. Third, admit a dimenssion to the gate only when its rule passes that audit. By this standard, our equity and robustness implementations are not ready to gate. That is not because these dimensions matter less, but because the rules we tried for them failed real patients. The admission table is a template the community can reuse to change that.

### What would change our conclusions

The claim would weaken if, at matched cost, the arithmetic mean accepted no more single-flaw tables than a calibrated gate on another cohort, or if calibrated gates failed their held-out audits elsewhere. It would also weaken if most real flaws turned out to be dense problems, each detectable on its own dimension, where sum-type rules are expected to gain [28]. Our dense test could not settle this, because two of its three components were below every rule’s detection limit. Each of these is testable with the protocol described here.

### Limitations

The evidence comes from one public cohort and one prediction task. Flaws are injected into real tables rather than produced by generators, whose failures may be denser. Bases, calibration runs and supplementary runs all reuse development patients in pseudo-roles. The calibration guarantee assumes a candidate is exchangeable with the fit runs, which holds by construction for the bases but must be argued for tables from a deployed generator. Privacy results are empirical lower bounds, not guarantees. The score mapping was fixed in advance, but its sensitivity analyses were post hoc, and they show that the size of the arithmetic mean’s excess depends on that choice. We did not measure how people rely on any of these rules. The protocol was amended twice, each time before the data concerned were scored. No result here constitutes clinical certification.

## 7 Conclusion

Evidence about synthetic clinical data is usually gathered carefully and then combined carelessly. When the same evidence was combined in different ways, across 50 independent real tables, averaging accepted tables with serious, known flaws that a gate rejected. The gate, in turn, was trustworthy only after it had been calibrated against real patients. Neither finding is complicated. What makes them useful is that both could be measured. The combining rule decides what passes, and it should be tested as carefully as the evidence it combines.

## Code and Data Availability

All code, configuration files and analysis scripts are available at https://anonymous.4open.science/r/calibrated-gate-artifact-2881/. The cohort is rebuilt from the public UCI Diabetes 130-US Hospitals dataset [9] by the included script.

## Ethics and Privacy Statement

This study uses a public, de-identified dataset under its repository terms and involves no human subjects. Its purpose is to make release decisions about synthetic patient data more protective of patients. One risk is that a calibrated gate is mistaken for a certification. We state throughout that verdicts are bounded by the evidence, the threat model and one cohort, and we report a case in which the privacy evidence itself misled.

## A Generators

All 12 generator tables failed the calibrated gate, each on both fidelity and utility. Privacy passed for Gaussian Copula and CT-GAN. It failed for TVAE and AIM, in both cases through the closer-than-real share (TVAE 19.6–22.2%), with the membership attack at chance. Each TVAE table also contained one or two training records, and one AIM table contained one; without the closer-than-real signal, the TVAE table with two copied records still fails privacy. TVAE’s synthetic outcome collapsed to a single class; the compensatory rules, which require every dimension score, did not score it.

The same twelve flaw–dose conditions applied to the three Gaussian Copula tables gave 36 flawed tables. Every rule, SDMetrics included, rejected all of them and the 12 unflawed baselines, so this set cannot separate the rules: no rule rescues a table whose generator has already failed.

## B Sensitivity Analyses

## C Calibration Diagnostics

On the 50 audit runs, the calibrated gate rejected 1, 2 and 3 runs at *α* = 0.025, 0.05 and 0.10 (nominal 1.0, 2.0 and 4.9 runs); privacy rejected none, and 2 conformal *p*-values fell at or below 0.05. Over 1,000 random fit/audit splits of the 100 original runs, refitted each time, the mean audit rejection rate was 1.98%, 3.92% and 9.81% (nominal 1.96%, 3.92% and 9.80%); at *α* = 0.05 the middle 90% of splits rejected 0–12% of their audit runs. Pooled with the 150 fresh runs, the gate rejected 2/200, 6/200 and 12/200 honest runs (one-sided 95% upper bounds 3.1%, 5.8% and 9.5%).

## D Implementation Details

*Signals*. Table 6 defines the *p* = 16 gating signals *h*_*j*_ (*X* ; *ρ*); retention counts once per probe and each calibration ratio twice, once per sign. The probes *q* are a random forest, an *l*_2_-regularised logistic regression and a gradient-boosting classifier; 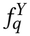 is probe *q* trained on table *Y* and AUC_*E*_ its AUC on *E*. For privacy, rows are encoded as standardised counts and one-hot categories. Bootstrap intervals are percentile intervals over 200 resamples of the rows of *E*, computed before orientation. Retention is undefined when the lower 95% limit of 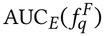 is at most 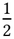, and the calibration ratios when the predictions on *E* are constant.

## GenAI Usage Disclosure

The authors used Claude (Anthropic) to help implement and test the experiment and analysis code, to write plotting scripts for the figures, to search for and check related literature, and to draft and revise the manuscript text. The authors ran all experiments on their own hardware. Every reported number was generated from saved run outputs by scripts included with the submission, and every citation was checked against its publisher or index record. The authors reviewed all content and take full responsibility for it.

